# Transient and sustained control mechanisms supporting novel instructed behavior

**DOI:** 10.1101/341818

**Authors:** Ana F. Palenciano, Carlos González-García, Juan E. Arco, María Ruz

## Abstract

The success of humans in novel environments is partially supported by our ability to implement new task procedures via instructions. This complex skill has been associated with the activity of control-related brain areas. Current models link fronto-parietal and a cingulo-opercular networks with transient and sustained modes of cognitive control, based on observations during repetitive task settings or rest (Dosenbach et al. 2008). The current study extends this dual model to novel instructed tasks. We employed a mixed design and an instruction-following task to extract phasic and tonic brain signals associated with the encoding and implementation of novel verbal rules. We also performed a representation similarity analysis to capture consistency in task-set encoding within trial epochs. Our findings show that both networks are involved while following novel instructions: transiently, during the implementation of the instruction, and in a sustained fashion, across novel trials blocks. Moreover, the multivariate results showed that task representations in the cingulo-opercular network were more stable than in the fronto-parietal one. Our data extend the dual model of cognitive control to novel demanding situations, highlighting the high flexibility of control-related regions in adopting different temporal profiles.

## Introduction

Following verbal instructions could seem, at first glance, a trivial aspect of human behavior, perhaps due to the easiness that we often experiment when following commands in our daily life. However, in continuously changing environments, the ability to use instructions to guide actions is essential for fit performance. In fact, this skill defines a crucial distinction between us and non-human apes: using language to share task procedures freed us from slow trial-and-error learning (Cole et al. 2013). Despite the biological relevance of this complex, flexible skill, some important aspects of its underlying neural architecture remain unknown. In the present study, we employed functional magnetic resonance imaging (fMRI) and both univariate and multivariate approaches to describe the transient and sustained control processes that allow us to follow novel verbal instructions.

The transformation of an instruction into effective behavior involves different processes. First, rules are semantically *encoded*, and proactive control processes (Meiran 1996; Braver 2012) are deployed to build a representation of the task (the so-called *task-set*; Sakai 2008). This set can be activated in advance (Meiran 2010; Ruge et al. 2013), biasing task-relevant processing in sensorimotor regions (e.g. Sakai and Passingham 2003; Sakai and Passingham 2006; Ekman et al. 2012; González-García et al. 2016; González-García et al. 2017) and thus, allowing us to *prepare*. Once the task context has been instantiated, task-sets must be *implemented* (Stocco et al. 2012), and reactive control processes become crucial (Cole et al. 2017), as they allow the inhibition of previously relevant action plans and the selection of target stimuli among possible distractors (Botvinick et al. 2001; Braver 2012). These proactive and reactive neural mechanisms, necessary for successful task encoding and implementation, have received considerable attention in the broader literature of cognitive control (e.g. Braver 2012; Palenciano et al. 2017).

Traditionally, the experimental approaches employed to study cognitive control use rather repetitive paradigms, which trigger proactive task-set reconfiguration with alternations between few rules (e.g., task switching; Monsell 2003) and/or reactive adjustments via conflict (e.g., the Stroop task; Stroop 1935). The evidence so far shows the involvement of a set of frontal and parietal areas during the execution of a wide spectrum of effortful, controlled tasks (Duncan 2010), including novel task execution (e.g., González-García et al. 2017). Due to the tight functional coupling of these regions (Fox et al. 2005; Seeley et al. 2007; Cole and Schneider 2007), they are often considered a unitary control brain network (namely, the Multiple Demand Network or MDN; Duncan 2010; Fedorenko et al. 2013). However, recent advances in experimental design and data analysis have led to its subdivision into at least two components-the cingulo-opercular and the fronto-parietal networks (CON and FPN, respectively)-, which seem to act at different, complementary time scales (Dosenbach et al. 2006; Dosenbach et al. 2008). The CON is comprised by regions that show both preparatory (cue-related) and sustained (across multiple trials) activations (Dosenbach et al. 2006), and has been associated with the proactive activation and maintenance of task-sets (Dosenbach et al. 2007). Conversely, FPN regions present mainly transient, cue and error-locked activity (Dosenbach et al. 2006) and their role has been described in terms of phasic, reactive adjustment of behavior (Dosenbach et al. 2007).

Support for this dual distinction comes not only from the analysis of sustained and transient neural signals while participants perform different tasks (Dosenbach et al. 2006), but it has also been confirmed when analyzing the information encoded in multivoxel activity patterns in those regions (Crittenden et al. 2016) and in functional connectivity data (both in resting state and on task; Dosenbach et al. 2007; Crittenden et al. 2016). Nevertheless, it has also been evidenced that such dual functioning, and specially the sustained involvement of the CON, is absent in certain task contexts (for example, when stimuli contain enough perceptual information to guide the response; Dubis et al. 2016). Last, crucially to the current study, it remains unknown whether there is a differential involvement of the two systems during goal-directed behavior in contexts of novelty. As novel tasks entail higher control demands than practiced ones (Norman and Shallice 1986), it is expected that they would be associated with a greater recruitment of maintained and transient processes mediated by CON and FPN, which could highlight their distinction.

Research in recent years has explored the brain regions underlying the encoding and implementation of instructions, and the specific roles carried out by each one (Brass et al. 2017). The findings so far support the involvement of the two main nodes of the FPN, the inferior frontal (IFS) and the intraparietal sulcus (IPS; e.g. Ruge and Wolfensteller 2010; Dumontheil et al. 2011; Muhle-Karbe et al. 2017), as expected from Dosenbach and colleagues’ model. The lateral prefrontal cortex (LPFC) in general, and the IFS in particular, have been linked to the encoding of new instructions (Hartstra et al. 2011; Demanet et al. 2016), showing higher activity in novel compared to practiced contexts (Cole et al. 2010; Ruge and Wolfensteller 2010). This region may be in charge, specifically, of the formation of novel stimulus-response mappings (when comparing against the formation of stimulus-stimulus associations; Hartstra et al. 2012). This supports its involvement in proactive processes related to the creation of novel task-sets, and not in the mere declarative maintenance of instructions in working memory (Hartstra et al. 2012; Brass et al. 2017). The IPS has shown, generally, a similar pattern (Ruge and Wolfensteller 2010; Dumontheil et al. 2011), although there is also evidence of a less abstract, sensorimotor representation in this region (Hartstra et al. 2012; Muhle-Karbe et al. 2014; González-García et al. 2017). Importantly, the functional coupling of the IFS and IPS with other brain regions contains fine-grained information about the content of novel instructions (Cole, Reynolds, et al. 2013). These distributed mechanisms of task-set representation also add evidence for the joint activation of fronto-parietal regions as a coherent functional system.

On the other hand, the CON network consists of the dorsal anterior cingulate (dACC), the anterior insula/frontal operculum area (aI/fO) and the anterior prefrontal cortex (aPFC). In contrast to the FPN, evidence of its involvement during instructed behavior is scarce. The dACC has been associated, in this context, with the reactive inhibition of irrelevant actions that interfere with the proper response (Botvinick et al. 2001; Brass et al. 2009). However, existing evidence does not yield strong support for a role of the dACC or the aI/fO in the encoding and/or maintenance of new instructed rules. The aPFC, in contrast, has been highlighted as a key region in the construction of novel task-sets, but only when rules are complex or abstract (Cole et al. 2010). Thus, the CON has not shown, as a system, a consistent behavior as the one predicted from the dual model framework.

The differential support for the participation of the two networks in novel instructed behavior could be due to different reasons. On the one hand, the nature of the behaviors explored could weight on transient mechanisms (FPN) to a higher extent than on sustained ones (CON), which besides of being more resource consuming (Braver 2012), develop in a time scale that may not be optimal in this context. In other words, the activity maintained in CON areas could be maximally beneficial when the relevant rules are stable in time (as in classic control paradigms), but not if quick task-set reconfigurations take place in a trial-by-trial fashion. In accordance with this idea, it has been proposed that reactive mechanisms are key to potentiate flexibility in novel instruction following (Cole et al. 2017). On the other hand, the evidence to date is scarce in contexts where novel instructions are embedded in designs aimed at isolating both control modes, which by definition act at different temporal scales.

When employing fMRI mixed designs (Petersen and Dubis 2012), the combination of events and blocks allows for the disambiguation of transient and sustained neural signals. To date, only one instructions study has been carried out using mixed designs (Dumontheil et al. 2011), and it employed complex practiced commands. These authors manipulated task-set complexity and studied transient activations linked to the encoding and implementation of instructions, while the sustained activations were analyzed only during implementation. Surprisingly, only two regions were involved in their sustained results: the IFS and the aPFC. Thus, the equal involvement of regions from both networks leaves open the role of the CON in instructed task execution and more importantly, whether this pattern applies to novel contexts.

We aimed to conduct an experiment which specifically tested the involvement of the dual control system proposed by Dosenbach and colleagues (Dosenbach et al. 2006; Dosenbach et al. 2008) during novel, instructed behavior. To do so, we adapted an instruction-following paradigm (González-García et al. 2017) to an fMRI mixed design, manipulating the experience with the instructions (novel vs. practiced) in different blocks of trials. This allowed comparing novelty-related activity patterns (i.e., sustained and phasic activations) against a control practiced condition. Furthermore, we aimed to better characterize the sustained activation profile associated with the CON. As the standard univariate analyses employed in previous studies did not help to clarify the information held by these networks, other plausible hypotheses in addition to proactive control involvement have been proposed (e.g., tonic attention maintenance; Coste and Kleinschmidt 2016). To address this issue, we employed recent multivariate techniques (Haynes and Rees 2006), an approach that has been shown to be highly informative. For example, using a combination of Multi-Voxel Pattern Analysis (MVPA) and Representational Similarity Analysis (RSA; Kriegeskorte et al. 2008), Qiao and colleagues (Qiao et al. 2017) were able to characterize how FPN areas adaptively change the task-set being represented, and how this process deals with interference from previous relevant rules. The dual-network model would predict a better maintenance through time of task-sets in CON, complementing the quick adjustment of the information encoded across the FPN. Thus, we employed RSA to assess whether the spatially distributed task representations were more consistent over time in CON than in FPN areas.

## Methods and materials

### Participants

37 students from the University of Granada, all right-handed and with normal or corrected-to-normal vision were recruited for the experiment (20 women, mean age = 21.13, SD = 2.47). All of them signed a consent form approved by the Ethics Committee of the University of Granada and received payment (20 to 25€, according to their performance) or course credits in exchange for their participation. Two participants were excluded from the final sample due to excess of head movement (> 3 mm). Sample size was selected according to recommendations for mixed designs (Petersen and Dubis 2012).

### Apparatus and stimuli

We used a total of 120 verbal instructions similar to those employed by González-García and colleagues (González-García et al. 2017). They were all composed by a condition and the two responses associated with the condition being true or false (e.g.: “*If there are four happy faces, press L. If not, press A*”). Half of the instructions referred to faces (their *gender* -female, male-, *emotional expression* -happy, sad-, or both), whereas the remaining referred to letters (their *type* -vowel, consonant-, *color* -blue, red-, or both). The instruction could also specify the *quantity* of specific stimuli, their *size*, or the *spatial contiguity* between them. Finally, the motor responses indicated a left or right index button press (“*press A*” or “*press L*”, respectively). Face and letter sets were equivalent in terms of these parameters. We conducted a pilot behavioral study to ensure that the difficulty was equivalent across the whole set. Then, to shorten task duration for the fMRI protocol, we built up six 100-instructions lists from the pool (again, equating face and letter-related elements) and assigned them to the participants, so each individual instruction was presented with the same frequency across our sample.

For each instruction, we built two grids of target stimuli: one fulfilling the condition specified (match) and the other one not (mismatch). They all consisted of *unique* combinations of 4 faces and 4 letters, which were drawn from a pool of 16 pictures: 8 face images (2 men and 2 women, 2 with happy expression and 2 sad, each in two different sizes -big/small-) from the Karolinska Directed Emotional Faces set (Lundqvist et al. 1998) and 8 letter images (2 consonants and 2 vowels, 2 in red color and 2 in blue, each in two different sizes -large/small-). Grids from face and letter instruction sets were built in parallel (establishing an equivalence between gender-letter type and emotion-color). Across the whole sample of participants, all instruction-stimuli (matching and mismatching grids) and instruction-response combinations (press A if true, press L if false; or the opposite) were employed.

The task was created with E-Prime 2.0 (Psychology Software Tools, Pittsburgh, PA). Inside the scanner, it was projected onto a screen visible through a mirror located on the head coil.

### Procedure

Participants performed a task in which they implemented novel and practiced verbal instructions referring to letters or faces, inside the fMRI scanner. The timing of the whole task was adapted to match the TR of the EPI sequence (2.21s), anchoring each event to the beginning of a scan acquisition, due to requirements of the FIR analyses conducted (see *fMRI analysis section*). Each trial (Fig. 1) started with the presentation of a verbal instruction (25.75°; *encoding phase*) during 2.21s (i.e., one TR), followed by a jittered interval with a fixation cross (2.21-8.84s, mean =5.525s). The grid of stimuli (21°) then appeared for 2.21s, where participants had to respond (*implementation phase*) using button boxes compatible with the scanner environment. The following trial began after a second jittered delay (with the same characteristics as the previous one).

**Figure 1:**
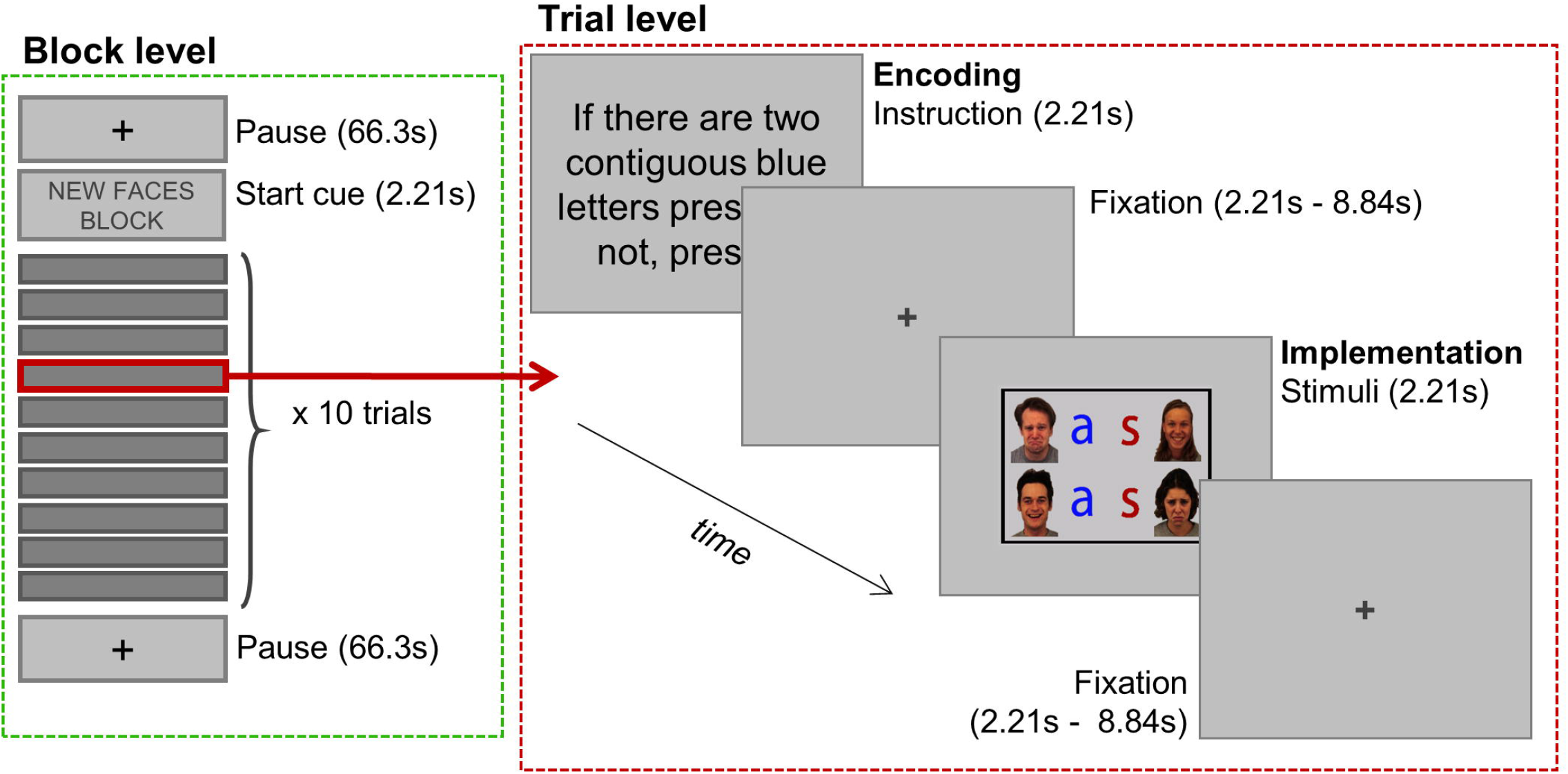
Mixed-design behavioral paradigm.

We were interested in two variables: the experience that the participants had with the trials (new vs. practiced) and the category of stimuli that the instructions referred to (faces vs. letters), having four possible conditions: Faces/New, Letters/New, Faces/Practiced, Letters/Practiced. As we employed a mixed fMRI design for our task, we manipulated those variables between blocks, for a total of 16 blocks (4 of each condition), with ten trials each. All blocks began with a cue indicating the experience and category condition (2.21s) followed by a jittered interval (2.21-8.84s, mean = 5.525s), after which the first trial began. Blocks lasted 154.7s, and were followed and preceded by pause periods of 66.3s (also indicated by pause cues of 2.21s). Importantly, pause duration was chosen to be long enough to ensure a robust baseline for block-related activity. The task was split into four runs, each composed of four blocks, one per condition. We carefully counterbalanced the order of blocks, ensuring that all of them were preceded and followed by the others the same number of times. Runs lasted 17.05 minutes, and the whole task 67.3 minutes.

Participants came to the laboratory approximately 24 hours before the fMRI session, and performed 10 repetitions of two blocks of ten instruction-grid pairings each (i.e., Faces/Practiced and Letter/Practiced blocks), which conformed the practiced instructions. Feedback was administered after each trial in this practice session, and learning was assessed in a pre-scanner test, with a requirement of at least 85% correct responses to continue the experiment. Across participants, all materials were equally employed in new and practiced conditions.

### FMRI: acquisition and analysis

MRI data was collected using a 3-Tesla Siemens Trio scanner at the Mind, Brain, and Behavior Research Center (CIMCYC, University of Granada, Spain). We used a T2*-weighted Echo Planar Imaging (EPI) sequence (TR = 2210ms, TE = 23ms, flip angle = 70º) to obtain the functional volumes. These consisted of 40 slices, obtained in descending order, with 2.3mm of thickness (gap = 20%, voxel size = 3mm^2^). The 4 runs consisted of 468 volumes each. We also acquired a high-resolution anatomical T1-weighted image (192 slices of 1mm, TR = 2500ms, TE = 3.69ms, flip angle = 7º, voxel size = 1mm^3^). Participants spent approximately 90 minutes inside the MRI scanner.

We used SPM12 (http://www.fil.ion.ucl.ac.uk/spm/software/spm12/) to preprocess and analyze the data. The first four volumes of each run were excluded to allow for stabilization of the signal. The remaining images were spatially realigned, time-corrected and normalized to the MNI space (transformation matrices were estimated from EPI images, and applied to them in the same step). Finally, they were smoothed using an 8mm FWHM Gaussian kernel. We built our experimental task on the basis of a mixed design (Petersen and Dubis 2012). Therefore, for each subject, we created a GLM including, simultaneously, events (separately, encoding and execution phases) and block regressors for each of the four conditions, to perform the main univariate analysis of this data. Events were modeled using a Finite Impulse Response (FIR) basis set (9 stick functions, encompassing 19.89s -9 TRs-following the onset of the events), while blocks were convolved with the canonical hemodynamic response (HRF) function (Visscher et al. 2003). We also modeled the pause periods (HRF convolved) and the block/pauses starting cues (FIR modeled), and included the errors (boxcar functions with same duration as the full trials, convolved with the HRF) and six movement parameters as nuisance regressors. A 756s high pass filter was set, taking into account block duration and the maximum time elapsed between events of the same condition.

At the within-subject level of analysis, we conducted *t*-tests comparing event regressors against the implicit baseline, time bin by time bin, separately for each condition. *T*-tests were also conducted to contrast blocks with pause periods (both collapsing across conditions, and separately), and also to compare between blocks of different conditions. At the group level, separate analyses were carried out for the sustained and transient components, in both cases correcting for multiple comparisons using a *P* <0.05 FWE cluster-wise criterion (from an initial uncorrected *P* < .001). In the first case, we used one sample *t*-tests with the subjects’ block contrast images obtained from the first level analyses. For the transient activity, we included the statistical maps obtained from the event contrasts into two ANOVA (encoding and implementation), performed as a full factorial design in SPM12 (Hartstra et al. 2011; Hartstra et al. 2012) and including Experience (novel, practiced), stimulus Category (faces, letters) and Time (9 time bins) as factors. This SPM design was chosen because it facilitates contrast specification, especially in complex models such as the one employed here. Nonetheless, all results were replicated with a repeated measures ANOVA also including a Subject factor, following an SPM flexible factorial model (Glascher and Gitelman 2008). We assessed main effects of experience and category, and their interaction with time bin. In the interaction of experience with time bin during the implementation stage, significant clusters were too big and extended over several different areas, so we adopted a stricter cluster forming threshold (uncorrected *P* < 0.001) to obtain smaller, anatomically more constrain clusters. Finally, to establish the directionality of these effects, we extracted the beta values of the significant clusters and compared the estimated hemodynamic response across conditions, both plotting the data, and performing post-hoc pairwise comparisons (Bonferroni corrected) with the SPSS software (SPSS 20.0 for Windows, SPSS, Armonk, NY).

We additionally performed non-parametric inference (based on 10.000 permutations and cluster-forming threshold of *P* < .001) on sustained activity data, using the software SnPM (http://www.sph.umich.edu/ni-stat/SnPM). We could not follow this strategy with the transient activity analysis, as the repeated-measures ANOVA design was too complex to implement with the software available. Nonetheless, it is noteworthy that the block non-parametric results successfully replicated the output from the parametric approach.

To further characterize these findings, we carried out three additional analyses. First, we performed a conjunction test (Nichols et al. 2005) to assess the overlap between areas showing sustained and transient (encoding and implementation) activity. To do so, we thresholded (*P* < .05 FWE cluster-wise criterion) the statistical maps obtained from the following contrasts of interest: (1) *t*-test of novel vs. practiced blocks, (2) main effect of Experience during the encoding of instructions; (3) and interaction of Experience*Time during the implementation stage. These three statistical maps were post-hoc selected based on the findings obtained from the analyses described above and our hypothesis regarding the roles of the CON and the FPN. These images, after being binarized, were used to assess the intersection of the contrasts. As a result, we obtained voxels significantly activated in all three situations simultaneously.

Next, we evaluated the congruency of our results with the proposal of Dosenbach and colleagues (Dosenbach et al. 2008) of two subnetworks for cognitive control. Specifically, we assessed the extent of overlap of the regions showing sustained and transient activations in our experiment with the CON and the FPN, respectively (Dosenbach et al. 2008). For this, we built spherical 10mm radius ROIs centered on the nodes of the CON (dACC [0, 31, 24], aPFC [-21, 43, -10; 21, 43, -10], aI/fO [-35, 18, 3; 35, 18, 3]), and FPN (IFS [-41, 23, 29; 41, 23, 29], IPS [-37, -56, 41; 37, -56, 41]), as published in Fedorenko and cols. (Fedorenko et al. 2013). ROI definition, including sphere size selection, was conducted following the parameters in the study of Dumontheil and colleagues (Dumontheil et al. 2011), in order to facilitate comparisons. The network templates were then overlaid against the thresholded statistical maps that we obtained in our results (using the same contrast images as in the conjunction analysis), after which we assessed which ROIs were present in each map and the percentage of voxels of each subnetwork involved in the different contrasts (Woolgar et al. 2016). It is important to note, however, the descriptive nature of our approach, as it did not involve the computation of inference statistics. This was due to the complexity of the mixed design analysis (which did not allow to obtain equivalent homogeneous statistics from both event and block-related signals). Nevertheless, the chosen procedure provided an informative comparison of the dual model (Dosenbach et al. 2008) and the sustained and transient activations estimated in our study.

Finally, we conducted a multivariate analysis to study the fine-grained distributed representation of instructions and their consistency along trial epochs (i.e., from the encoding to implementation stages). Specifically, we aimed to test differences in representation persistence between the two networks, and how novelty modulated this effect. To that end, we entered the non-normalized and unsmoothed functional images into a GLM similar to the specified above, with the exception that blocks were not defined and event regressors were convolved with the HRF. This modeling approach was selected because at this point there was no risk of misattributing the signal from transient and sustained components, and more importantly, because it provided a single parameter image for each event condition (instead of nine). The beta coefficient maps extracted (32 in total, corresponding to the encoding and implementation phases of each condition and run) were used to build a 32x32 Representational Dissimilarity Matrix (RDM; using The Decoding Toolbox; Hebart et al. 2014) for each FPN and CON ROI (as defined above), which had previously been inverse-normalized and coregistered to the participants’ native space. In the RDMs, each column and row corresponded to a different regressor, and each cell_i,j_ to the distance (computed as 1 - Pearson correlation) between the multivariate activity pattern associated with regressors i and j. Pearson correlation values were first normalized using Fisher’s z-transformation. We focused on the quadrant of the RDMs capturing the dissimilarities between encoding and implementation of instructions, in which the diagonal represented distances within different stages of same condition trials, and the off-diagonal represented values of different condition trials (Fig. 2). We computed the average difference between off and on-diagonal values for each ROI (González-García et al. 2018), as an index of representational consistency along time. Concretely, this index showed how similar the patterns of activations at the implementation and encoding stages of same condition were, in comparison with different condition trials. An index of 0 means that the information encoded in multivariate patterns was independent between encoding and implementation, while higher values reflect greater correspondence between the information encoded in both phases. We first checked that the index was significantly above 0 across regions using one-sample *t*-tests. As the aim of this analysis was to assess whether the consistency index varied between the FPN and the CON, we averaged the values of ROIs pertaining to each system and performed a paired *t*-test between them. Even when our main hypothesis-driven approach for this analysis was to group the regions into two segregated control networks (Crittenden et al. 2016), we also wanted to explore differences that could arise among areas of the same component -as there is no reason to assume that they all perform identical computations. To assess this possibility, we conducted a repeated-measures ANOVA within each network, with ROI as factor, which was later qualified with planned comparisons, Bonferroni-corrected. Finally, we obtained the consistency indexes separately for novel and practiced trials, and explored this effect with a repeated-measures ANOVA with Network and Experience as factors.

**Figure 2:**
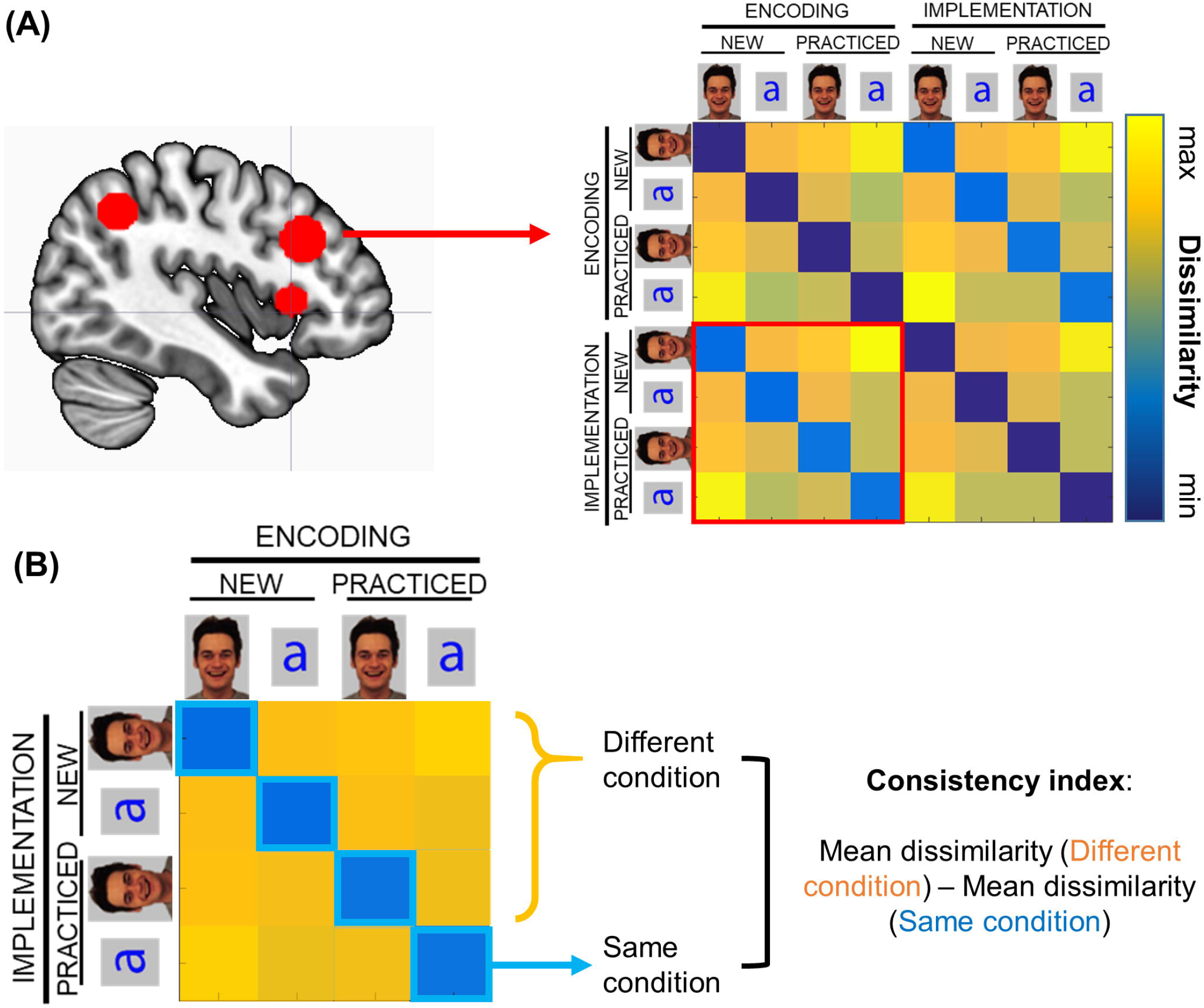
Representational Similarity Analysis. (A) First, a representational dissimilarity matrix (RDM) was built using the data of each cingulo-opercular and fronto-parietal region of interest. Each cell of the matrix indicates the dissimilarity between the representation of each pair of trial conditions at encoding and implementation stages. (B) The left lower quadrant was selected in each RDM. Within this quadrant, the diagonal (cells in blue) show dissimilarities between the encoding and the implementation of same-condition trials, and the off-diagonal values (cells in orange) refer to different-condition trials. Those values were averaged separately and subtracted to compute the persistence index employed in the analysis.

## Results

### Behavior

We analyzed the behavioral performance during the scanning session using two-way repeated-measures ANOVAs, with Experience (new vs. practiced) and Category (faces vs. letters) as factors. We found a significant effect of Experience in accuracy (*F*_1,34_ = 51.12, *P* < .001, η_*p*_^2^ = .601), with better performance for practiced (M = 94.7%, SD = 5.3) than for novel trials (M = 88.7%, SD = 6.8). The effect of Category was also significant (*F*_1,34_ = 5.31, *P* < .027, η_*p*_^2^ = .135), with better performance for faces (M = 92.6%, SD = 6.0) than for letters (M = 90.9%; SD = 7.4%). Finally, RT data from this session replicated the significant main effect of Experience (*F*_1,34_ = 290.48, *P* < .001, η_*p*_^2^ = .895), and also showed a significant interaction between Experience and Category (*F*_1,34_ = 32.56, *P* < .001, η_*p*_^2^ = .489), with faster responses to faces (M = 747.0ms, SD = 196.7ms) than letters (M = 783.6ms, SD = 188.7ms) in practiced trials, and the opposite pattern in novel ones (Faces: M = 1047.7ms, SD = 183.7ms; Letters: M = 982.7ms; SD = 172.1ms). Finally, we performed two additional ANOVAs on accuracy and RT data including Run as a factor, to rule out possible fatigue effects on our behavioral measures. Neither the main effect of Run (accuracy: *F*_3,102_ = 1.99, *P* < .120, η_*p*_^2^ = .055; RT: *F*_3,102_ = 2.11, *P* < .104, η_*p*_^2^ = .058) nor its interaction with Experience or Category were significant (*Fs* < 1.02, *Ps* > .100). This was further confirmed with a Bayesian repeated-measure ANOVA, in which both the main effect of Run and its interactions showed a BF_10_ < .3, strongly supporting a null effect of this variable and, thus, confirming that participants’ performance was stable across the whole task.

### fMRI

We first conducted a *univariate analysis* to assess sustained and transient activity, with the goal of exploring the effect of the experience with the task (new vs. practiced). As specified before, we also carried out a *multivariate analysis*, focused on the within-trial time scale, to study the consistency of multivoxel representation along phases of the task (encoding and implementation).

#### Univariate analysis

##### Transient activity

Event-locked activations were estimated using a set of FIR functions, obtaining nine parameters per regressor defined at the within-subject level. Then, they were entered into two separate ANOVAs: one to capture phasic activations associated with the encoding of instructions, and the other for their implementation. In both, we assessed the main effect of Experience, and its interaction with Time.

During the encoding of instructions (Table 1, Fig. 3), the main effect of Experience was significant bilaterally in the dorsolateral prefrontal cortex (DLPFC) -including the IFS-, and aPFC. To explore the directionality of this result, we extracted the beta estimates for each conditions and time bin (averaged across participants). Intriguingly, the hemodynamic response (HDR) was more pronounced for practiced compared to novel instructions in both DLPFC clusters (see Fig. 3). In the aPFC, beta values were also higher in the practiced condition, but in that case the HDR did not resemble the typical curve (see Fig. 3), but showed a deactivation, less pronounced for practiced rules.

**Table 1.** Transient activity results.

| **Label** | **ANOVA term** | **Direction** | **Peak coordinate** | **Z value** | **k** |
| --- | --- | --- | --- | --- | --- |
| *Encoding phase* |  |  |  |  |  |
| Left aPFC | Main effect | P > N | -36, 44, 8 | 5.44 | 155 |
| Left DLPFC | Main effect | P > N | -51, 20, 39 | 5.38 | 95 |
| Right aPFC | Main effect | P > N | 39, 47, 2 | 5.13 | 134 |
| Right DLPFC | Main effect | P > N | 48, 29, 29 | 4.64 | 91 |
| Cerebellum (lobule VI) | Interaction | N > P | -33, -43, -22 | 4.14 | 60 |
| *Implementation phase* |  |  |  |  |  |
| Left LPFC | Interaction | N > P | -45, 8, 29 | 7.61 | 430 |
| Right LPFC | Interaction | N > P | 54, 26, 26 | 7.12 | 295 |
| SMA/preSMA | Interaction | N > P | -3, 17, 53 | 6.86 | 177 |
| Right SPL | Interaction | N > P | 30, -55, 47 | 6.46 | 538 |
| Left SPL/IPL | Interaction | N > P | -24, -70, 44 | 6.11 | 516 |
| Right Fusiform gyrus | Interaction | N > P | 48, -58, -13 | 5.86 | 225 |
| Right aI/fO | Interaction | N > P | 33, 23, -4 | 5.75 | 112 |
| Left aI/fO | Interaction | N > P | -33, 23, -4 | 5.55 | 79 |
| Left Caudate | Interaction | P > N | -21, 8, 26 | 5.43 | 57 |
| Left SMG/STG | Interaction | P > N | -54, -37, 23 | 5.39 | 394 |
| Left BG / posterior insula | Interaction | N > P | -33, -19, -1 | 5.35 | 104 |
| Left fusiform gyrus | Interaction | N > P | -39, -46, -22 | 5.12 | 110 |
| Right SMG/STG | Interaction | P > N | 60, -34, 32 | 5.08 | 178 |
| Right BG / posterior insula | Interaction | - | 30, -19, 5 | 4.89 | 113 |
| Right MTG | Interaction | - | 48, -34, -10 | 4.73 | 48 |
| Left MTG | Interaction | - | -48, -22, -4 | 4.68 | 27 |
| Bilateral Caudate | Main effect | N > P | 3, 8, -4 | 4.47 | 106 |
| Right fusiform / PHG | Main effect | N > P | 27, -31, -22 | 4.11 | 75 |

Note: The ANOVA terms refer to the main effect of Experience and the interaction of Experience with Time (see *Methods* sections). The direction indicates whether the activity was higher in novel (N) or in practiced (P) conditions, while hyphens designate regions with no clear directionality (because the significant interaction term is driven not by heightened activation but by different timing of the response). Abbreviations stand for anterior prefrontal cortex (aPFC), dorsolateral prefrontal cortex (DLPFC), lateral prefrontal cortex (LPFC), supplementary motor area (SMA), presupplementary motor area (preSMA), superior parietal lobe (SPL), inferior parietal lobe (IPL), anterior insula/frontal operculum (aI/fO), supramarginal gyrus (SMG), superior temporal gyrus (STG), basal ganglia (BG), middle temporal gyrus (MTG), parahippocampal gyrus (PHG).

**Figure 3:**
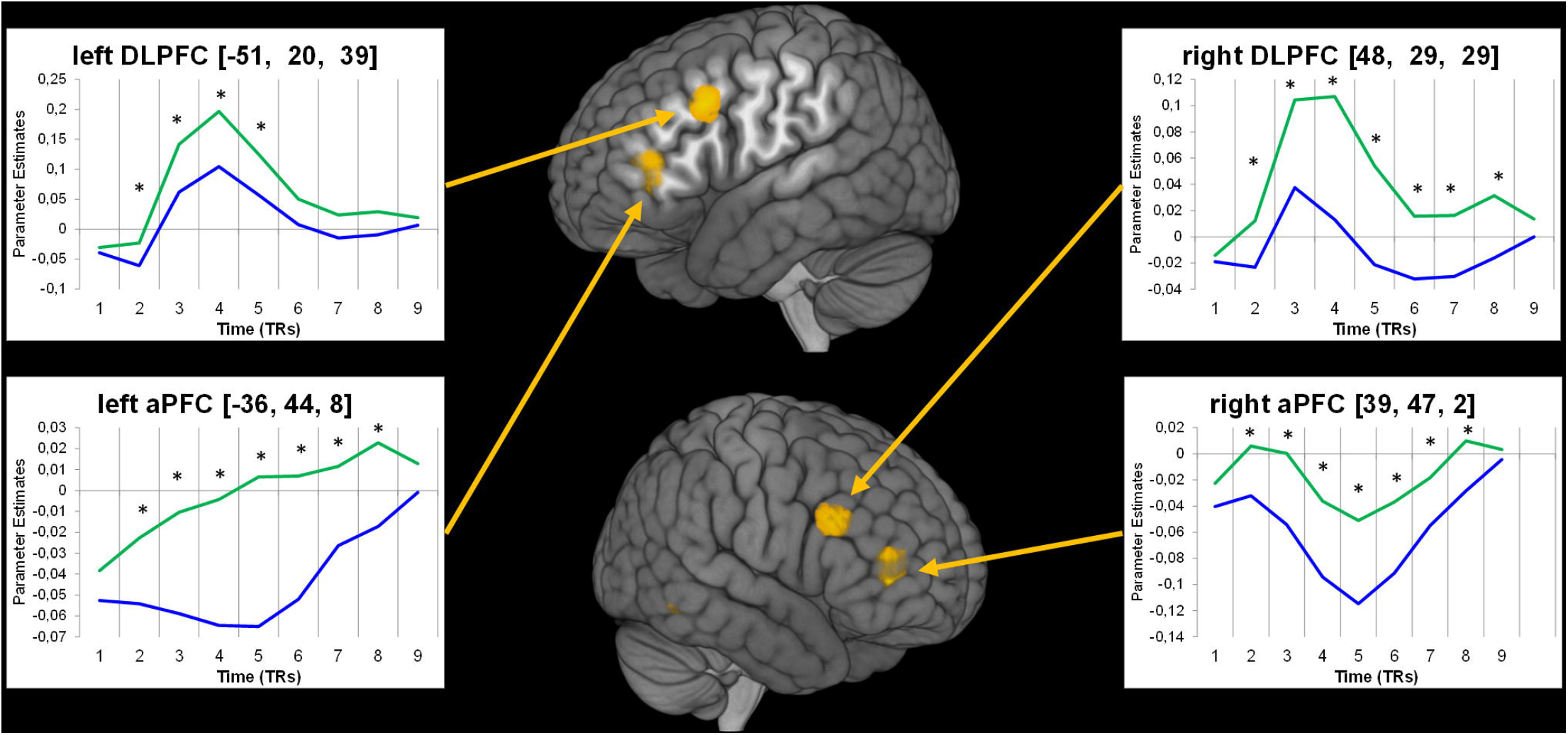
Results from the encoding stage ANOVA. Yellow clusters show regions where the main effect of Experience was significant. Insets show the hemodynamic response (beta values extraction) for novel (blue) and practiced (green) trials. Asterisks indicate that the conditions differed significantly (*P* < 0.05, Bonferroni corrected) in the corresponding time bin.

In contrast, a wide array of brain areas was differently activated in novel and practiced trials during the implementation of instructions (Table 1), as assessed by the interaction of Experience with Time (Fig. 4). As clusters were very large, we used a stricter statistical threshold to explore smaller, anatomically more accurate clusters (uncorrected cluster-defining threshold of *P* < .0001; this threshold was also employed to display the results in Fig. 4 and Table 1). In contrast to the encoding stage, almost all regions showed a higher HDR for novel than for practiced instructions, including the IFS, the inferior frontal junction (IFJ), the IPL and the aI/fO (Fig. 4). On the other hand, the bilateral supramarginal and superior temporal gyrus were more active in practiced trials

**Figure 4:**
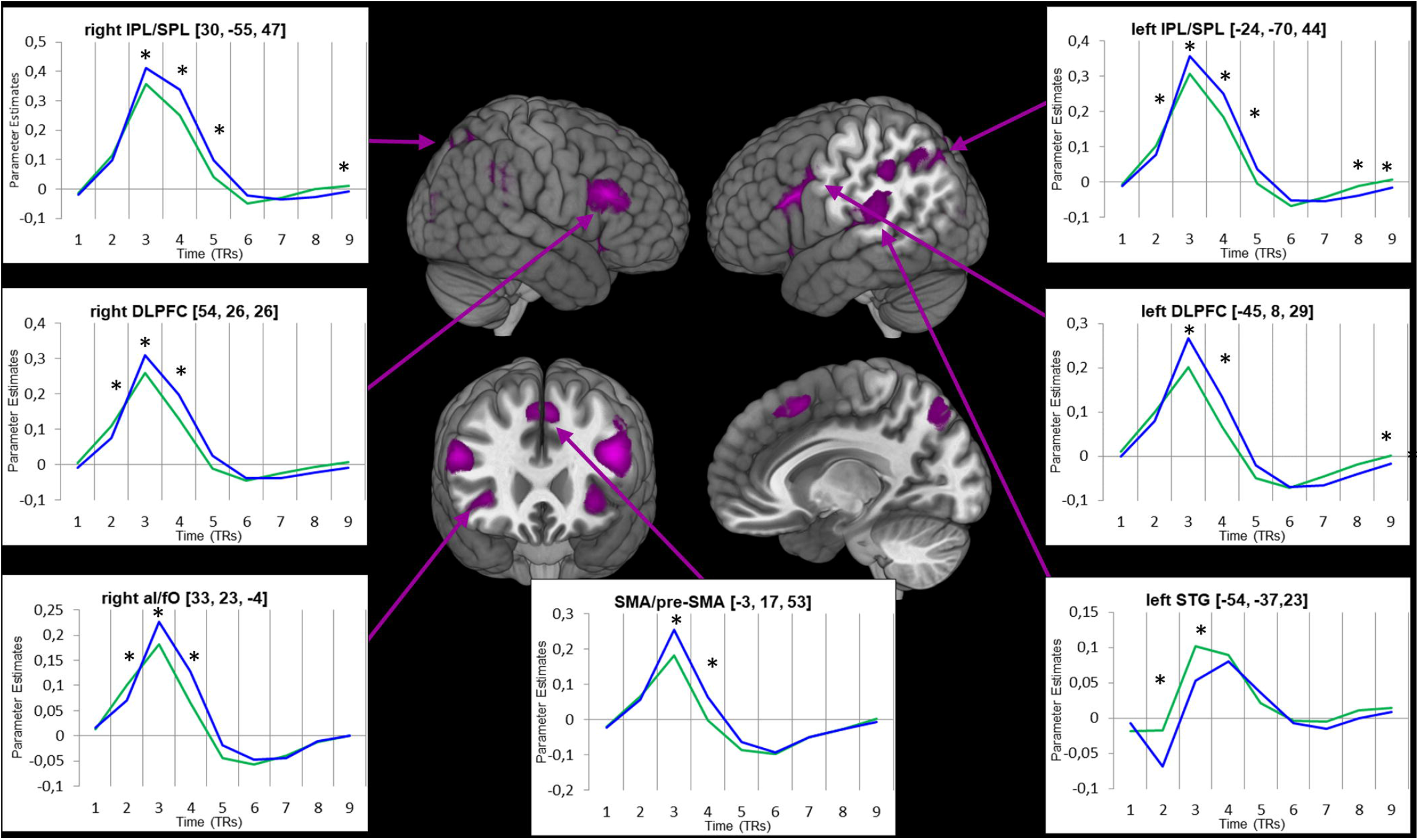
Results from the implementation stage ANOVA. Violet clusters show regions where the interaction of Experience and Time was significant. Insets show the hemodynamic response (beta values extraction) for novel (blue) and practiced (green) trials. Asterisks indicate that the conditions differed significantly (*P* < 0.05, Bonferroni corrected) in the corresponding time bin.

##### Sustained activity

We first aimed to detect areas showing sustained activity through long task blocks in comparison with rest, collapsing across all conditions. We did not observe any significant results in this analysis, nor when we compared just practiced blocks against baseline. On the other hand, sustained activity in novel blocks (vs. baseline) was found in the right aI/fO and bilaterally in the inferior parietal lobe (IPL), aPFC and DLPFC-also involving the IFS (Fig. 5A and Table 2). DLPFC and IPL were also significant when novel blocks were contrasted against practiced ones (Fig. 5B), providing support for their role for sustained control in new situations. Conversely, practiced blocks elicited higher sustained activity than novel ones in the ventromedial prefrontal cortex (vmPFC).

**Figure 5:**
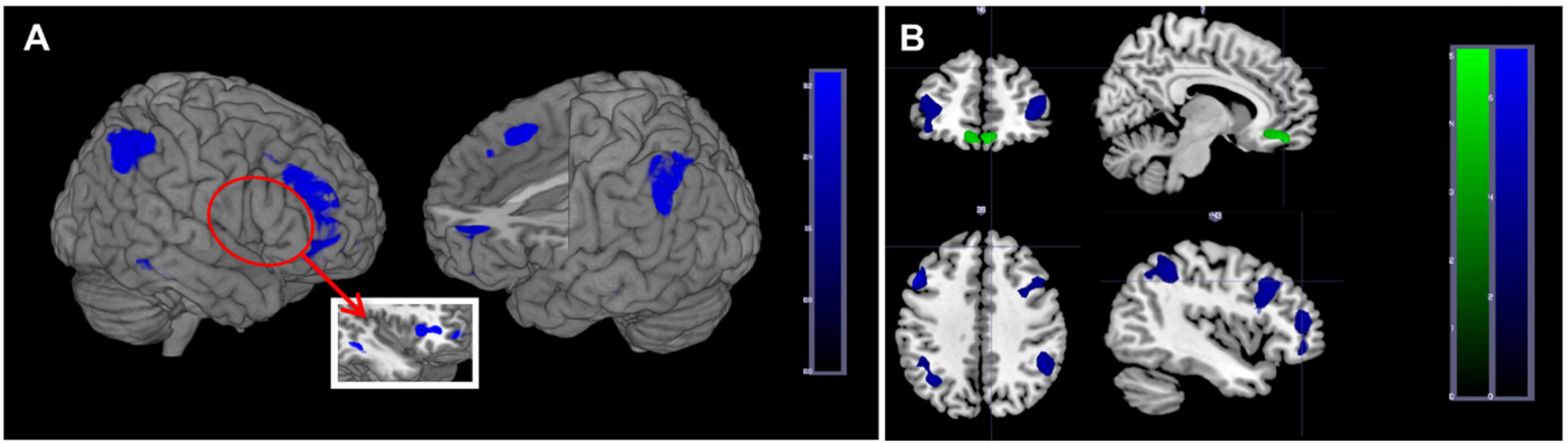
Sustained activity results. (A) Areas found in the *t*-test of Novel blocks against baseline. (B) Results from the contrast of novel versus practiced blocks. Clusters in blue show higher sustained activation in novel compared to practiced blocks, while the reverse is shown in green.

**Table 2.** Sustained activity results.

| **Label** | **Block labels** | **Peak coordinate** | **Z value** | **k** |
| --- | --- | --- | --- | --- |
| right IPL | Novel > Baseline | 45, -52, 41 | 6.03 | 340 |
| left IPL | Novel > Baseline | -42, -58, 47 | 5.27 | 330 |
| left MTG | Novel > Baseline | -54, -31, -10 | 5.48 | 122 |
| left aPFC/DLPFC | Novel > Baseline | -39, 47, 5 | 4.85 | 182 |
| right aPFC/DLPFC | Novel > Baseline | 39 53 -4 | 4.65 | 507 |
| bilateral SMA/preSMA | Novel > Baseline | -9 17 53 | 4.59 | 213 |
| right IFG/MTG | Novel > Baseline | 57, -25, -19 | 4.49 | 136 |
| right Cingulate gyrus | Novel > Baseline | 9, -28, 26 | 4.18 | 262 |
| left DLPFC/VLPFC | Novel > Practiced | -51, 20, 38 | 5.46 | 234 |
| right aPFC | Novel > Practiced | 39, 50, 5 | 4.37 | 142 |
| right IFJ | Novel > Practiced | 30, 11, 35 | 4.27 | 81 |
| left IPL | Novel > Practiced | -36, -61, 41 | 4.26 | 155 |
| right IPL | Novel > Practiced | 48, -49, 44 | 4.2 | 134 |
| left aPFC | Novel > Practiced | -45, 44, 14 | 4.16 | 143 |
| bilateral vmPFC | Practiced > Novel | 6, 47, -19 | 4.37 | 234 |

Note: Abbreviations stand for inferior parietal lobe (IPL), medial temporal gyrus (MTG), anterior prefrontal cortex (aPFC), dorsolateral prefrontal cortex (DLPFC), supplementary motor area (SMA), presupplementary motor area (preSMA), inferior frontal gyrus (IFG), middle frontal gyrus (MFG), lateral prefrontal cortex (LPFC), ventrolateral prefrontal cortex (VLPFC), inferior frontal junction (IFJ) and ventromedial prefrontal cortex (vmPFC).

#### Conjunction analysis

Results from our previous analyses suggested an overlap between regions with stronger sustained activity during novel blocks, and those with larger transient activity for the encoding of practiced instructions, and the implementation of novel ones. To quantify this observation, we performed an ad-hoc conjunction analysis with the corresponding three statistical maps obtained at the subject level (see *fMRI analysis* section). This test allowed us to confirm that one region, the left IFS, was involved across the three situations (Fig. 6).

**Figure 6:**
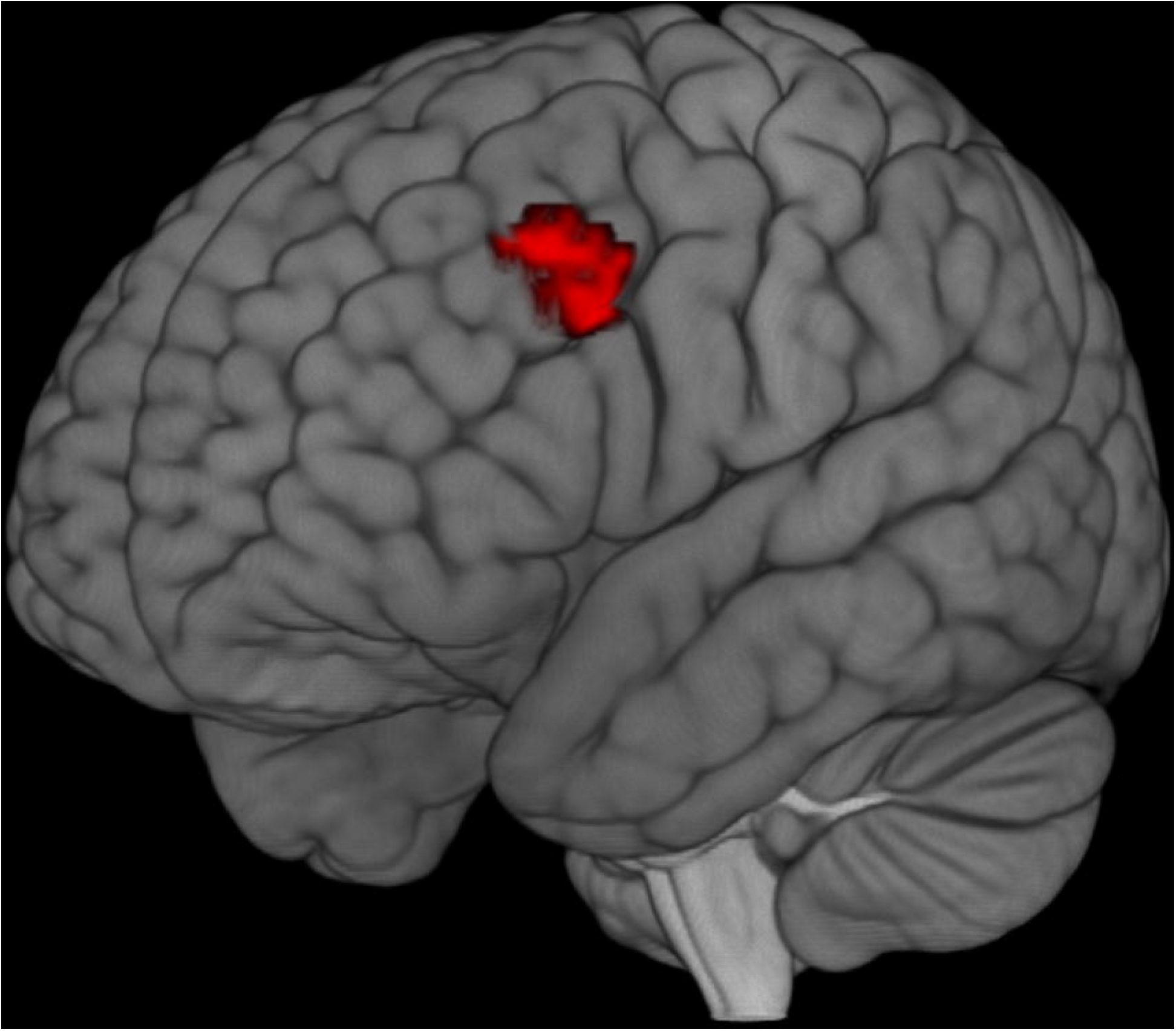
Results from the conjunction analysis. In red are voxels surviving to the conjunction test of (1) transient activity locked to practiced instructions encoding; (2) transient activity locked to novel instructions implementation and (3) sustained activity maintained through novel blocks. Peak coordinates: [-48, 20, 32], k = 63.

##### Network comparison

We also assessed the extent to which our principal sustained and transient results replicated previous findings regarding the involvement of two differentiable networks for cognitive control (Table 3): the CO and FP networks. Contrary to the framework put forward by Dosenbach and colleagues (Dosenbach et al. 2008), only the right aI/fO showed sustained activity throughout novel blocks, which just constituted 3.18% of the voxels of the CON template. Moreover, areas included in the FPN (bilateral IPS and the right IFS, involving a 42.92% of voxels of this network) were also present in the sustained activity maps.

**Table 3.**
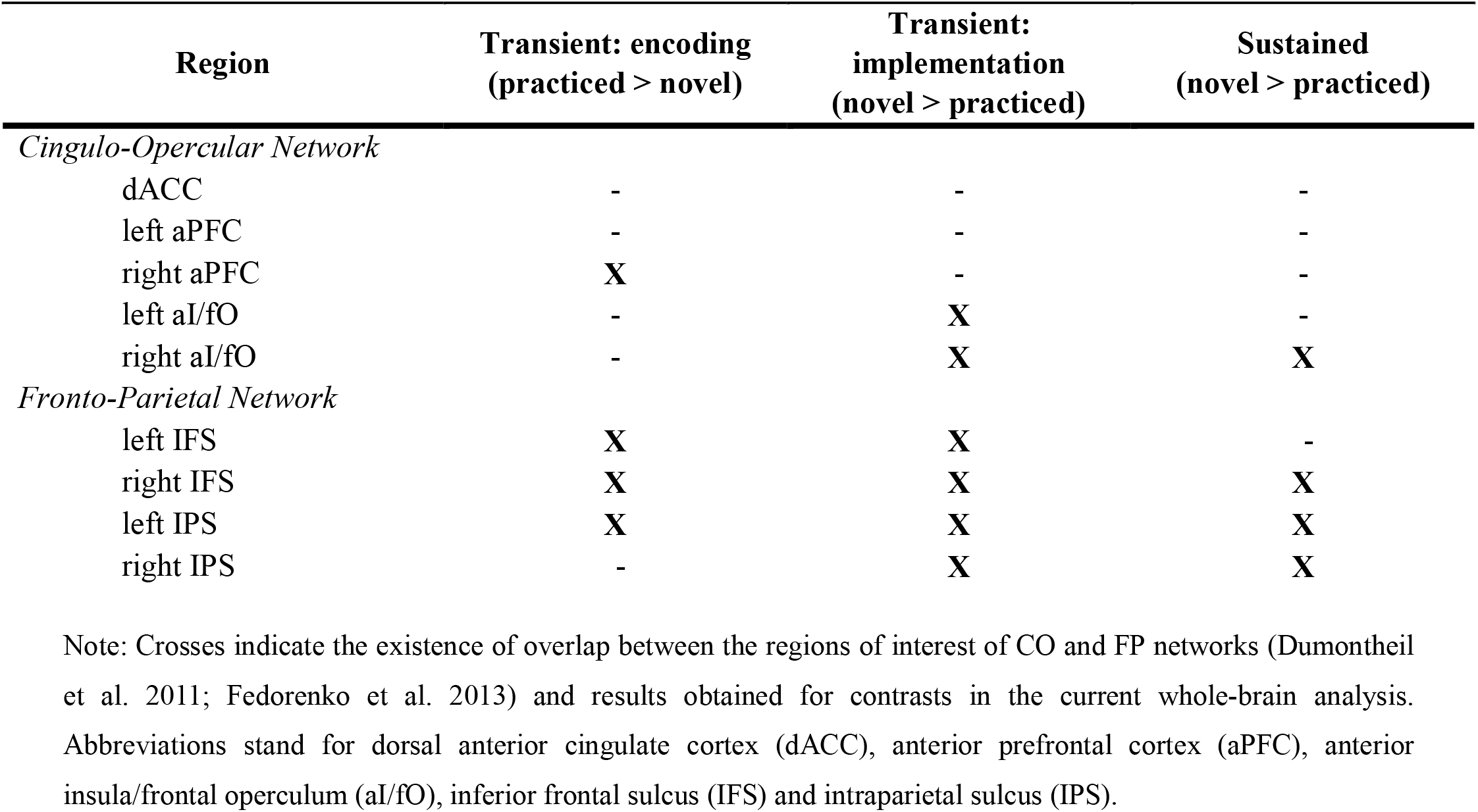
Transient and sustained signals at cingulo-opercular and fronto-parietal regions.

| **Region** | **Transient: encoding**<br>**(practiced > novel)** | **Transient:**<br>**implementation**<br>**(novel > practiced)** | **Sustained**<br>**(novel > practiced)** |
| --- | --- | --- | --- |
| *Cingulo-Opercular Network* |  |  |  |
| dACC | - | - | - |
| left aPFC | - | - | - |
| right aPFC | **X** | - | - |
| left aI/fO | - | **X** | - |
| right aI/fO | - | **X** | **X** |
| *Fronto-Parietal Network* |  |  |  |
| left IFS | **X** | **X** | - |
| right IFS | **X** | **X** | **X** |
| left IPS | **X** | **X** | **X** |
| right IPS | - | **X** | **X** |

Note: Crosses indicate the existence of overlap between the regions of interest of CO and FP networks (Dumontheil et al. 2011; Fedorenko et al. 2013) and results obtained for contrasts in the current whole-brain analysis. Abbreviations stand for dorsal anterior cingulate cortex (dACC), anterior prefrontal cortex (aPFC), anterior insula/frontal operculum (aI/fO), inferior frontal sulcus (IFS) and intraparietal sulcus (IPS).

At a transient time scale, the right aPFC, from the CON (4.69% of voxels), and the bilateral IFS and left IPS, from the FPN (18.61% of voxels), were involved during encoding of practiced instructions. During the implementation of novel ones, all ROIs of the FPN coincided with active clusters (although in an extent of just the 16.77% of the voxels), but were also accompanied by bilateral aI/fO from the CON (being, in this case, a 27.40% of CON voxels). Overall, the picture emerging from these comparisons is a mixture of CON and FPN involvement across both temporal modes of functioning.

##### Representational similarity analysis

In addition to the temporal profiles (transient vs. sustained) described above, differences between the CON and FPN may arise at a shorter time scale, within trial epochs. We explored this using RSA focused on the CON and FPN ROIs. We computed a consistency index associated with the maintenance of multivoxel representation of instructions from encoding to implementation stages (Qiao et al. 2017; see Fig. 2), in which larger values indicated a higher consistency along time (see *fMRI analysis* section). As expected, in all the regions examined, this index was significantly above 0 (all *P*s < .001 in one-sample *t-* tests) showing a correspondence between the information represented during novel instruction encoding and implementation. However, due to the temporal proximity of the source signal (consecutive events) this result could merely reflect the sluggish nature of BOLD response, although the jittered interval added between the encoding and the implementation should prevent or minimize this problem. In any case, this potential confound does not affect our analysis as we only focused in the relative differences in the index between both networks.

We first collapsed across novel and practiced trials, and observed that the CON’s consistency index was higher than the FPN’s one (*T*_34_ = 9.34, *P* < .001), suggesting more persistent task-set representations in the former network. We then explored variations within ROIs of both subnetworks, with two additional repeated-measures ANOVAs. In both systems, the effect of ROI was significant (CON: F_4,136_ = 91.84, *P* <.001, η_*p*_^2^ = .730; FPN: F_3,102_ = 30.64, *P* < .001, η_*p*_^2^ = .474) and planned comparisons showed that the differences were statistically significant between each pair of regions, except when they involved left and right portions of the same area. Within the CO subnetwork, the region showing the highest consistency over time was the bilateral aPFC (left: M = 1.028, SD = .207; right: M = 1.017, SD = .219), followed by the dACC (M = .850, SD = .207) and, finally, the aI/fO (left: M = .669, SD = .204; right: M = .672, SD = .171). On the other hand, the bilateral IFS (left: M = .821, SD = .231; right: M = .776, SD = .187) showed larger consistency than the IPS (left: M = .623, SD = .177; right: M = .583, SD = .151) in the FPN.

Finally, to assess whether this pattern was modulated by instruction novelty, we conducted an ANOVA with this variable and Network as factors. As expected, the main effect of Network was significant (*F*_1,34_ = 52.28, *P* < .001, η_*p*_^2^ = .606), and importantly, so was the main effect of experience with the task (*F*_1,34_ = 12.60, *P* = .001, η_*p*_^2^ = .270). Specifically, practiced instructions showed a higher consistency index than novel ones (novel: M = 0.745, SD = .195; practiced: M = 0.836, SD = .191), indicating that the experience facilitated a more efficient task-set maintenance within trials. The interaction term with Network was not significant, which suggests that the increase in similarity along the trial epochs with practice did not differ across CON and FPN regions.

## Discussion

In this study we investigated which brain networks underpin instruction following, and their fit within the dual control model (Dosenbach et al. 2006; Dosenbach et al. 2007; Dosenbach et al. 2008). To do so, we adapted a mixed design to a paradigm in which different novel and practiced instructions had to be encoded and implemented, and extracted the underlying transient and sustained brain signals. Our hypothesis was that novel instructions would recruit the CON and the FPN to a higher extent than practiced ones: the former proactively -transiently during instruction encoding, and in a sustained fashion across trials-, and the latter reactively -linked to the implementation stage-. Our results showed that the transient involvement of different regions varied depending on practice and the information stage (encoding vs. implementation) of instructions. Moreover, regions from both FPN and CON were involved both in the sustained maintenance of activity during novel blocks and during transient rule implementation. Multivariate patterns of activation in both networks showed a consistent differentiation between CON and FPN in how the information was maintained across the encoding and implementation stages, as the former network seems to hold instruction representations more consistently along time, an effect that increases with practice.

The analysis of transient activations by means of FIR models allowed to study how novelty influenced the regions engaged in a phasic mode during complex verbal instruction processing. In line with previous research (Ruge and Wolfensteller 2010; Dumontheil et al. 2011; Muhle-Karbe et al. 2017) we found that the IFS and the IPS, the main nodes of the FPN, were relevant at this time scale. Phasic activity was also found in the CON, concretely, in the aI/fO. In this sense, the whole pattern of regions presenting transient activity fits with our predictions based on Dosenbach’s model (Dosenbach et al. 2006). However, to better understand these findings, it is important to consider the two different processes that unfold along the trial epoch. We studied the encoding of instructions, more related with proactive preparation, and the subsequent implementation phase, where rules were applied to concrete stimuli, closely linked to reactive adjustments. During the initial encoding, no regions were transiently more active for novel than for practiced instructions. Conversely, the bilateral IFS was more active for practiced instructions than for novel ones. Later on, during the implementation, the IFS was again recruited, together with the IPS, the aI/fO and the preSMA. Importantly, here these regions showed larger activity for novel than practiced instructions, replicating previous findings (Ruge and Wolfensteller 2010; González-García et al. 2017).

The increased recruitment of the IFS in practiced compared novel instructions encoding may seem at odds with previous literature and our own predictions. Nonetheless, this finding may reflect the difficulty of fully preparing novel complex instructions during the encoding stage -in opposition with overly practiced ones, which could automatically retrieve the proceduralized task-set during this initial stage. In agreement with this, it has been previously proposed that novel rule preparation culminates when they are first implemented in behavior (Brass et al. 2009; Cole et al. 2013), an effect that may have been potentiated by the increased complexity and abstraction of our instructions in comparison with those used in previous research (e.g. Cole et al. 2010; Ruge and Wolfensteller 2010). As a result, the IFS activity may mediate practiced task-sets instauration and, as such, underlie a better proactive preparation in this condition. This is supported by the fact that this region has a relevant role in the preparation to implement instructions, in comparison with mere memorization demands (Demanet et al. 2016; Muhle-Karbe et al. 2017; Bourguignon et al. 2018).

Importantly, our conjunction results confirmed that the same left IFS cluster was present during the encoding of practiced instructions and the implementation of novel ones. Hence this region may underpin a preparatory process that can take place at different moments: earlier when the instruction is known (practiced) and its pragmatic representation can be retrieved, and later (i.e., when the stimuli are available) when we face a novel task, and this representation must be created from scratch. Nonetheless, which specific computations the IFS implements during this process is an open question. Different proposals have been made in the literature: binding of relevant stimuli and response parameters (Hartstra et al. 2012), mediating the transformation of semantic information into a pragmatic, action-oriented task representation (Ruge and Wolfensteller 2010), or maintaining the task-set in an active mode (Demanet et al. 2016), making it available for other lower-level regions. However, whereas novel instructions preparation seems to require the deployment of these three processes, practiced ones do not, as they do not need to be rebuilt but rather retrieved and updated. In light of our findings, therefore, task-set maintenance seems to be the most suitable common role underlying this region in both novel and practiced conditions. This is further supported by studies recording single and multiunit activity in monkeys’ LPFC (e.g., Freedman et al. 2001), which reveal the role of this area in the maintenance of different task-relevant information during delay periods.

Another remarkable set of results in the current study is the involvement of other regions during instruction implementation, such as the IPS and the preSMA. As implementation seems to rely to a high extent on reactive mechanisms, these regions may be implementing online control adjustments upon target presentation in novel trials, compensating for the less efficient proactive preparation during the encoding stage. From this perspective, the whole pattern of transient activations could be interpreted in terms of an interplay between proactive and reactive processes, which would depend on the novelty of the instructions that govern behavior. This interpretation fits with the balanced nature of proactive and reactive control modes: situations that weight proactive mechanisms to a higher extent trigger less reactive control, and vice-versa (Braver 2012). Nonetheless, it is also important to note that the temporal profile of activation of these brain areas is highly flexible. Whereas they have been linked to reactive functions (e.g., the preSMA seems to mediate the inhibition of irrelevant stimulus-response mappings in this context; Brass et al. 2009), patterns of activation consistent with proactive preparation have also been observed, such as increases of activity during encoding and preparation intervals (e.g. Hartstra et al. 2011; Dumontheil et al. 2011; Hartstra et al. 2012; Muhle-Karbe et al. 2014; Muhle-Karbe et al. 2017).

An additional core goal of our study was to extract sustained, block-wise activations to investigate whether a stable pattern of activation was maintained in CON and FPN areas during the execution of novel, demanding tasks, as it has been shown previously in more repetitive experimental settings (Dosenbach et al. 2006). In accordance with our expectations, blocks of new instructions were associated with a larger sustained recruitment of frontal and parietal regions, when comparing against both pause periods and practiced blocks. Nonetheless, the regions involved were more consistent with the main nodes of the FPN: the bilateral IFS and the IPS. Only the right aI/fO region and part of the aPFC, from the CON, showed sustained activation in novel blocks. Accordingly, when we explicitly tested the percentage of overlapping voxels between two networks and our results, we found higher coherence with the FPN. Our results aid to qualify the dual model of control, showing that sustained activation patterns are not the exclusive fingerprint of CON regions. In contexts of novelty, when higher flexibility is needed, nodes of the FPN are also recruited at this timescale, while sustained activity is restricted to certain nodes of the CON. This result may seem at odds with previous evidence. However, the nature of the behavior analyzed in our research departs considerably from the one captured by most of previous mixed-designed studies (Dosenbach et al. 2006), as our experiment required the continuous building and updating of novel complex task-sets. It has been argued that the sustained activation across the CON underlies the maintenance of relevant rules as long as they are needed (Dosenbach et al. 2008). While this mechanism may be efficient when the task remains the same, it may not be beneficial in long blocks where rules change in a trial-by-trial fashion. Here, the FPN may implement sustained control processes independent of the specific task-set adopted on each trial. Due to the role of this network in establishing the widest and most flexible pattern of connectivity with other brain regions (Cole et al. 2013), one possibility is that sustained activity across FPN regions implements some kind of tonic state of high efficiency in information routing between domain-specific regions. This view is supported by two different sources of evidence. First, task-dependent variability in the sustained engagement of CON has been previously reported, as in the case of perceptually driven tasks (Dubis et al. 2016). Second, sustained activity in lateral prefrontal and parietal cortices has also been found in studies which also relied on task-set updating: during blocks in which task switching was required (Marini et al. 2016), and while executing distinct instructions (Dumontheil et al. 2011). Overall, our findings highlight that both control networks, especially FPN areas, display a rather general ability to switch between phasic and tonic temporal modes depending on the nature of the tasks to be accomplished.

The result of our conjunction test, in which we identified common clusters at both phasic and tonic timescales, gains again relevance at this point. The same left IFS cluster involved transiently during the encoding of practiced instructions and the implementation of novel ones, which we propose underlie the maintenance of instructed task-sets, is also recruited in a sustained fashion through novel blocks. The relationship between the functions carried out at the two timescales is not straightforward; nonetheless, it is unlikely that they coincide, as this may result in an unnecessary redundancy across both timescales. It could well be the case that this and other regions perform distinct computations depending on temporal parameters, as previous neuroimaging data show that the LPFC, in general, can adopt different temporal dynamics (Jimura et al. 2010; Braver 2012). Results of the current investigation indicate that a demanding and rich task environment can recruit both temporal modes of functioning of this area, and moreover, that this profile is sensitive to the novelty of the situation. On the one hand, this evidence highlights the flexible nature of this brain region. On the other hand, such results could reflect an organizational principle by which different cognitive computations are multiplexed in distinct temporal dynamics within brain areas.

Finally, we also explored multivoxel activity patterns in both networks’ nodes, obtaining results consistent with the classic dual network model (Dosenbach et al. 2008). Areas within the CON represented task-sets more consistently over trial epochs, i.e., from encoding to implementation stages. This result strongly supports the proposal that these regions are in charge of maintaining information in a sustained, proactive fashion even in the absence of maintained univariate activation. Moreover, we found that this effect was affected by the experience with the trial: when the instructions were practiced, the consistency of the representation was higher, suggesting a possible mechanism by which the task representation gains in fidelity as it is repeatedly used. Interestingly, a recent study showed that task rule representation is more stable across the pre-target epoch when the instruction must be memorized in comparison with novel to-be-implemented ones (Muhle-Karbe et al. 2017). Overall, these results agree with the idea that novel trials require the semantic information of the instruction to be transformed into an action-related representation, a process that needs time to unfold and evolves up to target presentation. Moreover, this could explain why less reactive adjustments may be deployed when practiced instructions are translated into actions, as our results of transient activity during the implementation show.

Further research is needed to connect the scarce findings provided from this and other mixed design studies, and the broader cognitive control literature. For example, a recent study showed, employing MVPA, that task-sets were better encoded (i.e., decoded with higher accuracy) in FPN than in CON regions (Crittenden et al. 2016). These findings are not incompatible with ours, as we used RSA and our analysis was focused in the transference of rule representation between two temporal time points -and not in classification accuracies at concrete time points of the task. Nonetheless, due to the decision of using a mixed design to extract transient and sustained activations, our experiment was not optimized for performing MVPA on our data. Previous research (González-García et al. 2017) has shown that regions consistent with both CON and FPN encode the relevant stimuli category of the instructions, before its implementation. Future studies will help to characterize, from this approach, which information is contained in transient and sustained activation patterns -and whether this is segregated between the two control networks. Finally, it is important to highlight that the extent of novelty entailed by each instruction was limited, given that the global task structure remained the same throughout the experiment. To study control mechanisms acting in novel contexts, we generated a large amount of trials including unique task rules and complex and also unique target combinations (Cole et al. 2010; Hartstra et al. 2011; González-García et al. 2017). However, target categories (faces and letters) and motor responses (employing the two index fingers) remained the same across the whole task. While fixing these parameters allowed us to exert experimental control, the complexity of novel situations that humans face daily is far richer and more variable. Future studies should aim for increasingly more ecological paradigms, where the general task structure also varies in a trial-wise fashion.

## Conclusions

The current study provides insights about the dual network perspective of cognitive control, expanding this model to novel complex task contexts. Crucially, results indicate that even when the two networks are functionally differentiated, both seem act at both tonic and phasic timescales during novel instruction processing. Furthermore, the division between proactive and reactive control does not seem to be mapped in a straightforward way into these two networks. Future studies must be conducted to further detail their contributions. Specifically, the computations and information held at the sustained time scale remain unknown, as also their relationship with mechanisms that develop at a faster, transient scale. The expansion of multivariate decoding techniques could help to better disentangle between the computational roles of both neural networks.

## Acknowledgments

This work was supported by the Spanish Ministry of Science and Innovation (PSI2016-78236-P to M.R.) and the Spanish Education, Culture and Sport Ministry (FPU2014/04271 to A.F.P.). This research is part of A.F.P.’s activities for the Psychology Graduate Program of the University of Granada in Spain. We are grateful to Ben Inglis (https://practicalfmri.blogspot.com/) for his kind help during the implementation and testing of the MRI acquisition sequences employed in the current experiment.

